# Extraction of naringin dihydrochalcone and its effect on reducing blood lipid levels in vitro

**DOI:** 10.1101/2024.01.28.577672

**Authors:** Mu-yang Tian, Si-qi Li, Jing-wen Qi, Lei Zhang, Xiao-lei Yu

## Abstract

Response surface methodology (RSM) was used to optimize the extraction process of naringin. The central component design included three parameters of extraction, namely, temperature (X_1_), solid-liquid ratio (X_2_), and extraction time (X_3_). The optimum extraction temperature was 67℃; the ratio of material to liquid was 54:1mL/g, and the extraction time was 2.8h. According to the best extraction conditions, naringin was processed to verify the accuracy of the model. Five parallel experiments were set up, and a yield of 32.48 mg/g naringin was obtained, that was equivalent to the predicted yield of 32.56 mg/g. Naringin was purified to obtain naringin-refined products using DM101 macroporous adsorption resin. Naringin dihydrochalcone was synthesized following catalytic hydrogenation of purified naringin. The structures of naringin and naringin dihydrochalcone were determined via Fourier infrared spectrometer and nuclear magnetic resonance spectrometry. Evaluate the ability of naringin dihydrochalcone to bind with glycine bile sodium and bovine bile sodium by simulating the gastrointestinal environment in vitro. Further focusing on HepG2 cells, a high cholesterol induced high-fat HepG2 cell model was established. We measured the effects of different concentrations of naringin dihydrochalcone on intracellular lipids in denatured HepG2 cells and further validated the lipid-lowering effect of naringin at the cellular level. The results showed that naringin dihydrochalcone has potential application in functional foods for lowering blood lipids.

## 1 Introduction

With the improvement of modern living standards, people’s dietary structure and lifestyle habits are also changing, and the incidence of hyperlipidemia is constantly increasing. The clinical manifestation of hyperlipidemia is that the plasma lipid content in the patient’s body is significantly higher than the normal level, which is a cardiovascular system disease that poses great harm to the human body[1]. Pomelo is the mature fruit of the citrus plant in the Rutaceae family, and is one of the main fruit resources in China. The outer skin of pomelo is relatively thick, accounting for about 30% to 60% of the total amount of pomelo. Whether consumed fresh or in the production of fruit juice and jam, a large amount of by-products such as skin and residue are produced, and the utilization rate of these by-products is extremely low. 90% of them are discarded as garbage, which seriously wastes resources and also causes great environmental pollution[2–3]. TANG et al. It has been found that naringin dihydrochalcone has a certain effect on diabetes[4]. In addition, the effective group of naringin dihydrochalcone with antioxidant effect - 2,6-dihydroxyacetophenone structure is realized by eliminating peroxide and hydroxyl radical[5]. At present, studies on the hypolipidemic effect of naringin dihydrochalcone are limited. A previous study by our the research group showed that naringin exerts a blood lipid-lowering effect [6]. In this study, naringin monomer was extracted from naringin and catalysed hydrogenation to prepare naringin dihydrochalcone. The ability of naringin dihydrochalcone to bind sodium glycine cholate and sodium bovine cholate was evaluated by simulating gastrointestinal environment in vitro, so as to evaluate its lipid-lowering activity. HepG2 cells, a human liver cancer cell line, are commonly used by medical researchers worldwide to study liver lipid metabolism. Therefore, this study further focused on HepG2 cells and established a high-fat HepG2 cell model induced by high cholesterol. The effects of different concentrations of naringin dihydrochalcone on the intracellular lipids of denatured HepG2 cells were measured, and the lipid-lowering effect of naringin dihydrochalcone was further validated at the cellular level, in order to provide theoretical basis for the development of naringin dihydrochalcone functional food.

## 2 Materials and Methods

### 2.1 Materials and Reagents

Pomelo peel was obtained from Rongxian County, Guangxi Province, China and served as a source of naringin. Naringin (purity > 98%) was purchased from Hefei Bomei Biotechnology Co. Ltd. (Anhui, China).

Sodium glycolic acid (purity > 98%) was purchased from Hefei Qiansheng Biotechnology Co. Ltd. (Anhui, China).

D101 macroporous resin, sodium taurocholate, Trypsin, Pepsin were acquired from Solebo Biotechnology Co., Ltd. (Beijing, China).

Chromatography-grade methanol was purchased from Tianjin Guangfu Reagent Co., Ltd. software (Tianjin, China).

Naringin dihydrochalcone standard (purity > 98%) was purchased from Xi’an Huilin Biological Products Co. Ltd. (Shanxi, China).

Palladium carbon purchased from Tianjin Baima Technology Co. Ltd. (Tianjin, China).

Spectral grade deuteroacetone was purchased from Shanghai Yien Chemical Technology Co. Ltd. (Shanghai, China).

Spectral grade potassium bromide was purchased from Tianjin Hengchuang Lida Technology Development Co. Ltd. software (Tianjin, China).

Spectral grade tetramethylsilane was purchased from Hangzhou Sloan Material Technology Co. Ltd. (Zhejiang, China).

HepG2 cells were purchased from the Shanghai Institute of Biology, Chinese Academy of Sciences. (Shanghai, China).

Ethanol, sodium hydroxide, sulfuric acid, hydrochloric acid, and other chemicals and solvents were of analytical grade and were purchased from Tianjin Tianli Chemical Reagent Co. Ltd. software (Tianjin, China).

Cholesterol (purity > 99%) was purchased from Sigma Chemical Co.St.(Louis,MO, USA).

Total cholesterol (Cat. No. A111-2-1),Triglyceride (Cat.No. A110-1-1), Low-density lipoprotein cholesterol(Cat. No. A113-1-1), and High-density lipoprotein cholesterol (Cat. No. A112-1-1) were obtained from the Nanjing Jiancheng Bioengineering Institute (Nanjing, China).

### 2.2 Optimization of the naringin extraction process

#### 2.2.1 Pomelo peel pretreatment

Wash the fresh pomelo peel, remove black spots, cut it into small pieces, and dry the pomelo peel pieces using an air blower dryer at 50 °C until the weight no longer decreases. The dried pomelo peel pieces are crushed into powder using a high-speed multifunctional grinder, filtered through a 60-mesh sieve to obtain coarse pomelo peel powder, dried and stored for future use[7–8].

#### 2.2.2 Establishment of a naringin standard curve

A 80-mg sample of naringin standard was accurately weighed and diluted with absolute ethanol in a 500-mL volumetric flask to obtain a 0.16 mg/mL naringin standard solution. Different volumes of the solution were then distributed into 10-mL colorimetric tubes (0, 0.3, 0.6, 0.9, 1.2, 1.5, and 1.8 mL). Next, 5 mL of 90% volume fraction ethylene glycol solution and 0.1 mL of 4 mol/L NaOH solution, add distilled water to 10 mL, and invert repeatedly until the solution is evenly mixed[9]. React in a 40 ℃ water bath for 10 minutes and measure absorbance at 420 nm. (UV Visible Spectrophotometer, UV-6300, Meipeda Instrument Co., Ltd, Shanghai, China). The average values of three replicates for each concentration were obtained. These data were then used to construct a naringin standard curve[10].

#### 2.2.3 Calculation of extraction rate of naringin

Weigh 5g of pomelo peel powder, set it to the corresponding temperature and stir continuously. After reaching the time, filter it with a vacuum pump and dilute the resulting filtrate to a 500 mL volumetric flask[11]. Measure the absorbance value according to the section 2.2.2, corresponding to the standard curve of naringin, calculate the concentration of the diluent, calculate the extraction rate according to the following formula, set three equilibria, and take the average value[12].

Naringin extraction rate (mg/g) = CV/W

Where: C - concentration of naringin in diluent (mg/mL);

V - volume of extract (mL);

W - mass of pomelo peel powder (g).

#### 2.2.4 Single factor experimental design

Naringin powder (0.5g) was accurately weighed, and the effects of different solid–liquid ratios, extraction times, and extraction temperatures on its extraction rate were investigated to determine the behaviour of its surface under the influence of each factor. An ethanol volume fraction of 75%, material-liquid ratio of 45:1 mL/g, extraction time of 2.0 h, and extraction temperature of 60 ℃ were used as the basic conditions for the single factor screening test. When studying a given factor, all the other conditions were kept constant [13].

#### 2.2.5 Response surface experiment design

On the basis of single factor experiment, according to the principle of Box-Behnken center combination design, the optimal extraction range was selected, with naringin concentration as the response value, extraction temperature (X_1_), solid-liquid ratio (X_2_) and extraction time(X_3_) as independent variables, and the extraction rate of naringin as the response value[14]. Data analysis was carried out using Desigan-Expert 8.0.6 software, and an optimization model of three factors and three levels was obtained, totaling 17 groups of experiments[15]. See Table 1 for the factors and level design of response surface test.

**Table 1.**
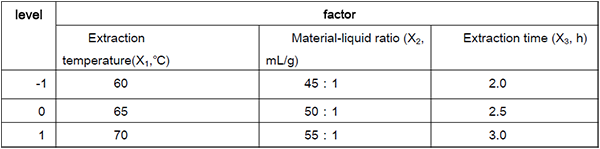
Response surface test design.

### 2.3 Purification of naringin

Naringin was separated and purified by DM101 macroporous resin, and the purification conditions were determined to be a loading concentration of 0.075mg/mL, a sample solution pH of 3.5, and a loading flow rate was 1.5 mL/min[16].

### 2.4 Structural identification of naringin

The Nicolet iS5 Fourier transform infrared spectrometer was used for sample infrared spectroscopy (FT-IR) detection. Potassium bromide tablet method is used for sample preparation. The sample is mixed evenly with potassium bromide and ground and pressed in an agate mortar[17–19].The scanning range is from 500 cm^-1^ to 4000 cm^-1.^ AVANCE III nuclear magnetic resonance spectrometer is used to detect the nuclear magnetic resonance hydrogen spectrum (1H NMR) and carbon spectrum (13C NMR) of the sample. Solvent: deuterated acetone, internal standard: tetramethylsilane (TMS) [20–22].

### 2.5 Preparation of naringin dihydrochalcone

Purified naringin (5.0 g) was placed in a stainless steel high-pressure reactor. Thereafter, NaOH solution was slowly added to obtain a reaction solution with pH 14. Next, excess 10% palladium carbon catalyst was also added and the pressure and temperature of the reactor were maintained at 1.6 MPa and 50 ℃, respectively. The reactor was then placed on a magnetic stirrer and hydrogen gas was introduced for 3 h before the reaction commenced. After 1 h, the hydrogen valve was closed to maintain the reaction rate. After 12 h, when the reaction reached completion, the product was filtered to recover the catalyst. Then, the filtrate was removed from the NaOH solution using 732 cation exchange resin, and the hydrogenation product was obtained after standing, filtering, and drying [23].

### 2.6 Structural identification of naringin dihydrochalcone

The method used was the same as that described in section 2.4.

### 2.7 In vitro study of the effects of naringin dihydrochalcone on lowering blood lipid

#### 2.7.1 Drawing the standard curve of cholic acid salt

Standard solutions of sodium glycocholic acid and sodium taurocholate at concentrations of 0.05, 0.10, 0.15, 0.20, and 0 were prepared. Thereafter, 2 mL of standard solution and place them in different stoppered test tubes[24]. Add 8 mL of sulfuric acid solution with a mass fraction of 60%, mix well, heat in an 80 ℃ water bath for 10 minutes, and quickly ice bath for 5 minutes. Then, measure the absorbance at a wavelength of 387 nm using UV visible spectrophotometry. Use absorbance as the vertical axis and bile acid content as the horizontal axis to draw the standard curves of glycine bile acid and taurocholate[25].

#### 2.7.2 Naringin dihydrochalcone binding cholate experiment

Naringin dihydrochalcone extracts (2 mL) at concentrations of 100, 200, 300, 400, and 500 mg/L, were each placed in a 100-mL triangular flask. Add 3 mL of pepsin solution (10 mg/mL) and 1 mL of HCl solution (0.01 mol/L) to each triangular flask, shake in a constant temperature shaker at 37 ℃ (simulating gastric digestion environment), and adjust the pH to 6.3 with NaOH solution (0.1 mol/L). Subsequently, 4 mL of trypsin solution (10 mg/m L) was added, and a constant temperature oscillator was used to oscillate for 1 hour at 37 ℃ (simulating intestinal environment). 4 mL of sodium glycyrrhetinic acid (0.4 mmol/L) was added to a sample flask; Add 4 mL of taurocholate (0.5 mmol/L) to another sample in a triangular flask. Shake in a constant temperature oscillator at 37 ℃ for 1 hour, centrifuge at 4000 r/min for 20 minutes, take the supernatant, and measure the absorbance value at 387 nm using colorimetric method. Each sample is measured three times in parallel, and the remaining content of glycylcholic acid and taurocholate is calculated according to the standard curve. The ratio of the difference between the total amount of glycylcholic acid or taurocholate added and the remaining amount to the total amount is the binding rate, expressed as a percentage. [26].

#### 2.7.3 Determination of conjugation rate of cholic acid salts

The ability of naringin dihydrochalcone to lower blood lipid level was determined based on the binding rate of the two sodium cholates with naringin dihydrochalcone [27].

#### 2.7.4 MTT cell proliferation and toxicity experiments

We used the MTT colorimetric method to detect the effects of different concentrations of naringin dihydrochalcone on the proliferation of Hep G2 cells. [28] For cell digestion (1–2 min), 0.25% trypsin was added to Hep G2 cells at the logarithmic growth stage cultured in MEM medium containing 1% dual antibody and 10% foetal bovine serum at 37 ℃ in a 5%-CO_2_ incubator. Thereafter, the cells were counted and inoculated into a 96-well plate (100 cells per well) and incubated at 37 ℃ in a 5%-CO_2_ atmosphere for 24 h until the cell count reach 5000 cells per well.

New culture media with the naringin dihydrochalcone at different concentrations (6.25, 12.50, 25.00, 50.00, 100.00, 200.00, 400.00, 800.00, and 1600.00 μg/mL) were added to the cell culture followed by further incubation for 24 h. Thereafter, the supernatant was discarded and the residue was incubated with 5 mg/mL MTT solution for 4 h, after which 100 μL of dimethyl sulfoxide (DMSO) solution was added. Further, after the crystal violet staining at the bottom of the incubator disappeared completely, the cells were inoculated into a 96-well plate on an enzyme-linked immunosorbent assay (ELISA) reader and absorbance measurements were performed at 490 nm. The formula for calculating cell survival rate was as follows:

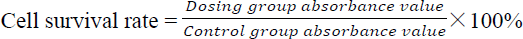

#### 2.7.5 High-cholesterol cell model

When Hep G2 liver cancer cells covered 80–90% of the bottom of the cell incubation bottle, the cells were digested with tripsin, suspended, and counted. Thereafter, the cells at a concentration of 2 × 10^5^ cells/mL were inoculated into a 96-well plate and incubated in a 5%-CO_2_ incubator for 24-48 h.

When the cell monolayer reached the bottom of the well plate, the culture medium was replaced with serum-free medium and cholesterol solutions of different mass concentrations (10, 15, 20, 25, and 30 μg/mL) were added. After incubation for 24 h, the culture medium was discarded and the cells were wash three times with phosphate buffered saline (PBS). The cells were further digested using trypsin and the cell suspension obtained was transferred into a centrifuge tube and centrifugation was performed at 2000 r/min for 6 min. After this step, the supernatant was discarded and the residue was washed with PBS. Centrifugation was further performed and the resulting supernatant was discarded. The cell residue was then resuspend in PBS and crushed using a cell crusher. The concentration of cholesterol in each cell group was then determined in accordance with the instructions provided with the reagent kit [29].

#### 2.7.6 Effects of naringin dihydrochalcone on TG, TC, LDL-C, and HDL-C levels in Hep G2 cells

When Hep G2 cells in the cell culture bottle covered 80–90% of the bottom of the cbottle, the cells were digested with trypsin, centrifuged, and diluted to a given concentration. This was followed by inoculation into a 96-well plate, the addition of 2.5 mL cholesterol at 10, 20, and 40 μg/mL to the complete culture medium, and incubation for 24 h. The supernatant was aspirated and the residue was digested using trypsin, centrifuged, and washed with PBS three times. Finally, 250 μL of 1% TritonX-100 was added followed by cracking in an ice bath at 4 ℃ for 40–50 min and the detection of TG, TC, LDL-C, and HDL-C levels according to instructions stated in the reagent kits [30].

### 2.8 Data processing

The test data were analysed using Design-Expert 8.0.6 and Origin 2021 software.

## 3. Results and Discussion

### 3.1 Determination of optimum extraction conditions of naringin

#### 3.1.1 Naringin standard curve

The standard curve of naringin is shown (Supplementary Materials Figure S1).

#### 3.1.2 Single factor experimental results

The effect of extraction temperature on naringin extraction rate is shown in Figure 1. From this figure, it is evident that with an increase in temperature, molecular movement intensified, resulting in the dissolution of naringin and a continuous increase in its extraction rate, which reached a maximum at 65 ℃. When the temperature was increased further, the naringin structure was destroyed and degraded; thus, the extraction rate decreased significantly. Further, Figure 2 shows the effect of the material-liquid ratio on the naringin extraction rate. From this figure, it is evident that an increase in material-liquid ratio, i.e., the extraction solvent amount, resulted in a rapid increase in the naringin extraction rate. Specifically, the highest extraction rate was observed when the material-liquid ratio was 50:1 mL/g. A further increase in the extraction solvent amount resulted in a rapid decrease in the naringin extraction rate. This observation can be explained as follows. With an increase in solvent extraction amount, the number of free water molecules and dissolved naringin molecules increased. However, with an excessive extraction solvent amount, more impurities were introduced, resulting in the observed decrease in the naringin extraction rate. Therefore, the optimal material-liquid ratio was 50:1 mL/g. As shown in Figure 3, with an increase in extraction time, the extraction rate of naringin increased gradually, and peaked at 2.5 h, after which it dropped sharply. This maximum extraction rate corresponded to when the material came into full contact with the solution. However, as the extraction time increased further, the viscosity of the material increased, resulting in a decrease in the naringin extraction rate. Therefore, 2.5 h was identified as the optimal extraction time.

**Figure 1.**
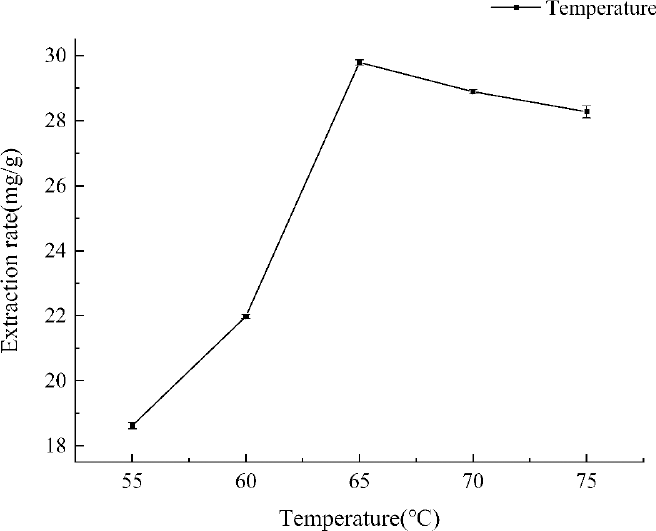
The effect of extraction temperature on the extraction rate of naringin.

**Figure 2.**
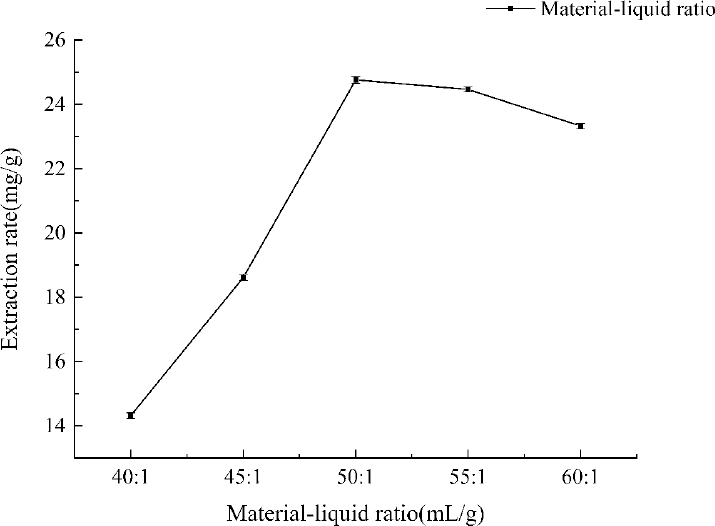
The effect of material-liquid ratio on the extraction rate of naringin.

**Figure 3.**
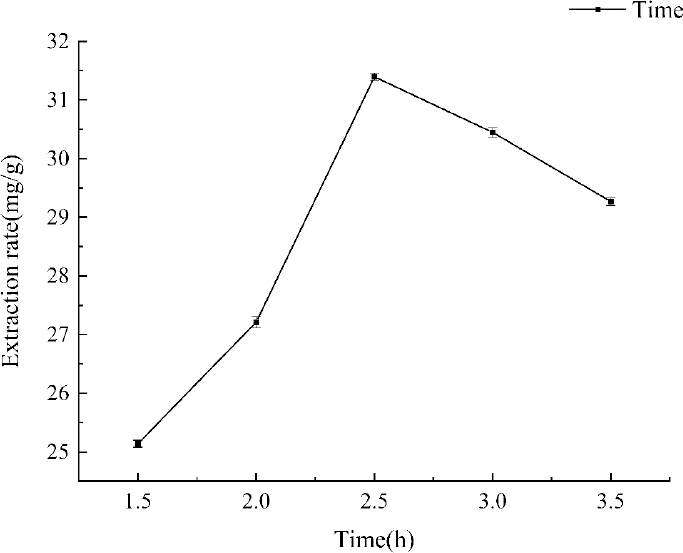
The effect of extraction time on the extraction rate of naringin.

#### 3.1.3 Response surface optimization test results and analysis of variance

The process and results of the response surface optimisation tests are listed in Table 2, while the variance analysis of the response surface model is presented in Table 3. Using Design-Expert 8.0.6 software, model fitting analysis was performed on the test results shown in Table 2, considering the extraction temperature (X_1_), material–liquid ratio (X_2_), and extraction time (X_3_) as independent variables, and the extraction rate (Y) as the response value. Thus, the following regression equation was obtained:

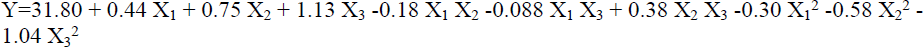

Via the analysis of variance (ANOVA), we further determined the quality of the model and clarified the relationship between various parameters. The misfit error obtained(P = 0.105 > 0.05) as shown in Table 3 indicated a good data fitting effect, with the primary and secondary terms reaching a highly significant level, and the interaction terms AB and BC showing significance. Further, the model design was reasonable, and the response surface model could accurately reflect the relationship between naringin yield and the extraction conditions. The accuracy test for the regression equation also showed that the coefficient of determination, R^2^ of the equation model, was 0.9948, which is indicative of the highly significant performance of the model. Furthermore, R^2^adj = 0.9881 could explain the 98.81% response value variation in the experiment, indicating that the experimental model had good fit with real data and has practical guidance significance. Therefore, the model can be used to analyse and predict the optimal naringin extraction process.

**Table 2.** Response surface optimization test results.

| Test number | $X_1$ | $X_2$ | $X_3$ | Extraction rate<br>mg/g |
| --- | --- | --- | --- | --- |
| 1 | 0 | 0 | 0 | 31.71 |
| 2 | 0 | -1 | 1 | 30.15 |
| 3 | -1 | -1 | 0 | 29.45 |
| 4 | 0 | 0 | 0 | 31.74 |
| 5 | -1 | 0 | 1 | 31.37 |
| 6 | 0 | 0 | 0 | 31.91 |
| 7 | -1 | 1 | 0 | 31.30 |
| 8 | 0 | -1 | -1 | 28.70 |
| 9 | 1 | 0 | 1 | 31.86 |
| 10 | -1 | 0 | -1 | 28.88 |
| 11 | 1 | 1 | 0 | 32.02 |
| 12 | 1 | -1 | 0 | 30.90 |
| 13 | 0 | 0 | 0 | 31.88 |
| 14 | 0 | 0 | 0 | 31.77 |
| 15 | 1 | 0 | -1 | 29.72 |
| 16 | 0 | 1 | -1 | 29.45 |
| 17 | 0 | 1 | 1 | 32.41 |

**Table 3.** Response surface optimisation test results.

| Source | Sum of Squares | Degrees of Freedom | Mean Square | $f$ Value | $p$ Value | Significance |
| --- | --- | --- | --- | --- | --- | --- |
| Model | 23.89 | 9 | 2.65 | 148.02 | < 0.0001 | ** |
| $X_1$ | 1.53 | 1 | 1.53 | 85.39 | < 0.0001 | ** |
| $X_2$ | 4.47 | 1 | 4.47 | 249.27 | < 0.0001 | ** |
| $X_3$ | 10.22 | 1 | 10.22 | 569.64 | < 0.0001 | ** |
| $X_1 X_2$ | 0.13 | 1 | 0.13 | 7.43 | 0.0295 | * |
| $X_1 X_3$ | 0.031 | 1 | 0.031 | 1.71 | 0.2326 | |
| $X_2 X_3$ | 0.57 | 1 | 0.57 | 31.79 | 0.0008 | ** |
| $X_1^2$ | 0.38 | 1 | 0.38 | 21.45 | 0.0024 | ** |
| $X_2^2$ | 1.43 | 1 | 1.43 | 79.60 | < 0.0001 | ** |
| $X_3^2$ | 4.57 | 1 | 4.57 | 255.05 | < 0.0001 | ** |
| residual | 0.13 | 7 | 0.018 |  |  |  |
| Spurious term | 0.094 | 3 | 0.031 | 4.05 | 0.1050 |  |
| Error term | 0.031 | 4 | 0.00777 |  |  |  |
| the sum | 24.02 | 16 |  |  |  |  |

Note: \* indicates significant difference, $P < 0.05$ \*\* Indicates that the difference is extremely significant, $P < 0.01$

#### 3.1.4 Response surface graphic analysis

The response surface analysis of the extraction temperature, liquid-to-material ratio, and extraction time is shown in Figures 4–9. Observing the steepness of the response surfaces using a 3D graph showed a steeper slope and a higher degree of inclination, indicating that the experimental results shown on the vertical axis were more sensitive to the factors shown on the corresponding horizontal axis, and that the interaction between the two factors had a stronger impact on the experimental results. From the perspective of the contour map transformed from the 3D graph, a saddle shaped contour indicated a significant interaction. The smaller the eccentricity of the ellipse, and the closer it was to a circle, the less significant was the interaction. Further, the steeper the response surface curve, the more significant was the impact. These criteria have been widely applied in other studies. Figures 4–9 also validated the ANOVA results. Among the three factors, extraction time showed the most significant impact, followed by the liquid to material ratio and extraction temperature. Further, we also found that among the pairwise interaction surfaces of the various factors, the longitudinal span of the interaction between the liquid to material ratio and extraction time was the largest.

Furthermore, the eccentricity of the contour ellipse was the largest. This indicated that the interaction between the two factors had the most significant impact on the naringin extraction rate. Additionally, the longitudinal span of the interaction between extraction temperature and the liquid to material ratio was the second largest, and similarly, the eccentricity of the contour ellipse was the second largest.

Our results also indicated that the longitudinal span of the interaction between extraction temperature and extraction time was the smallest, and the eccentricity of the corresponding contour ellipse was also the smallest. We also noted that the trend of effect of the interactions between extraction temperature, liquid-solid ratio, and extraction time on naringin extraction rate was: BC>AB>AC, consistent with the results of ANOVA. This also indicated that the model can be used to analyse and predict the optimal naringin extraction process conditions.

**Figure 4.**
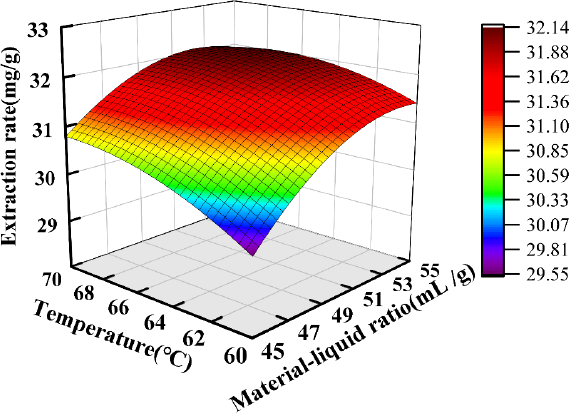
Material–liquid ratio and temperature response surface diagram.

**Figure 5.**
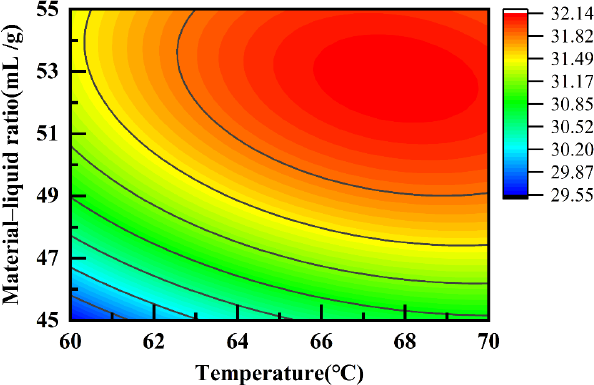
Temperature and material–liquid ratio interaction contour.

**Figure 6.**
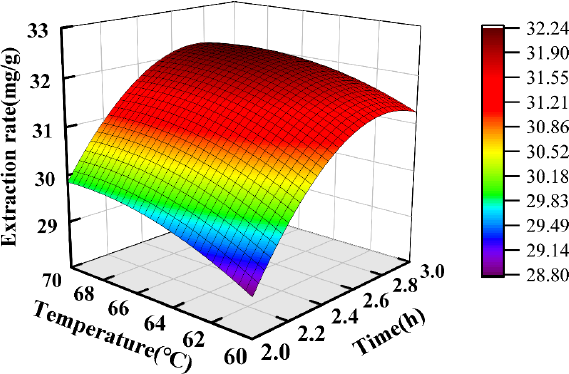
Temperature and time response surface diagram.

**Figure 7.**
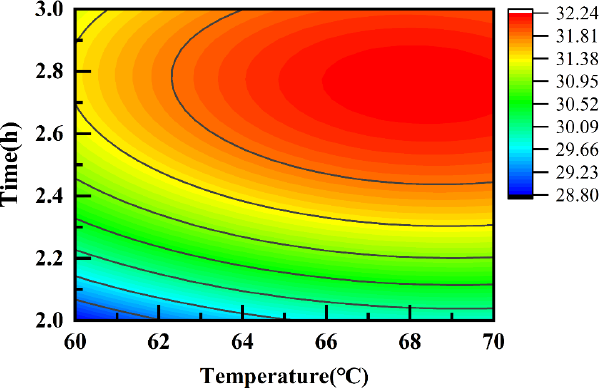
Temperature and time interaction contour.

**Figure 8.**
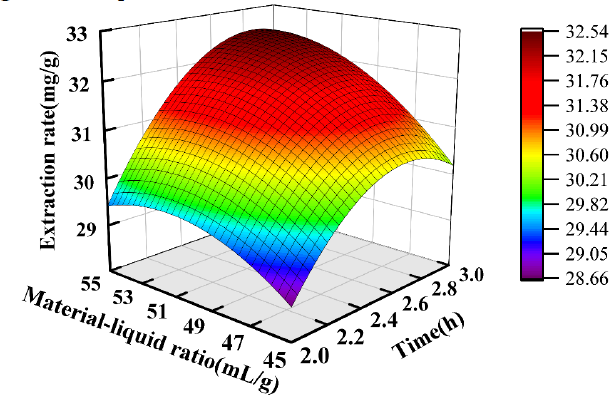
Material–liquid ratio and time response surface diagram.

**Figure 9.**
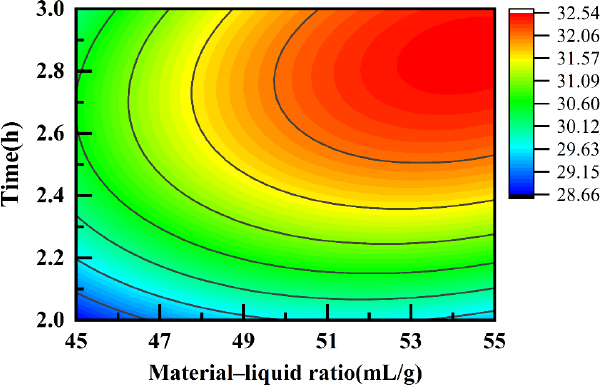
Material–liquid ratio and time interaction contour.

#### 3.1.5 Determination of optimal extraction conditions

Based on the response surface of each factor, it could be concluded from the highest point of the response surface that there exist extreme values for each factor within the set experimental range. According to the regression model prediction, the optimal naringin extraction conditions were determined as follows: temperature, 66.93 ℃; material–liquid ratio, 54:1 mL/g; and extraction time, 2.8 h. Under these conditions, the predicted yield of naringin was 32.56 mg/g. Further, considering the actual situation, the optimal process conditions were adjusted to a temperature, material–liquid ratio, and extraction time of 67℃, 54:1mL /g, and 2.8 h, respectively. The raw materials were processed under these optimal extraction conditions to verify the accuracy of the model. To this end, a total of five parallel experiments were set up, and the average yield of naringin was 32.48 mg/g. Thus, the extraction and determination model showed high accuracy, consistent with the predicted value. This indicated that the extraction method was feasible.

### 3.2 Structural analysis of naringin

#### 3.2.1 IR spectrum analysis

The IR analysis results for our refined naringin products are shown (Supplementary Materials Figure S2).

#### 3.2.2 NMR spectrum analysis

The NMR analysis of our refined naringin products are shown (Supplementary Materials Figure S3-6).

### 3.3 Structural analysis of naringin dihydrochalcone

### 3.3.1 IR spectrum analysis

The IR analysis results for our refined naringin products are shown in Figure 10. The strong and wide absorption peaks at 3000–3680 cm^-1^ were attributed to the alcohol and phenolic hydroxyl groups in naringin dihydrochalcone. The absorption peaks at 2840–2980 cm^-1^ were attributed to the C-H bond stretching vibration on the saturated carbon in the structure. Further, the peak at 1630 cm^-1^ was attributed to the carbonyl stretching vibration absorption peak in naringin dihydrochalcone, while the peaks at 1520, and 1440, and 1390 cm^-1^ were attributed to the C=C stretching vibrations of the aromatic ring in the structure. The multiple absorption peaks at 1200–1060 cm^-1^ were attributed to the C-O stretching vibrations of aromatic ether and fatty ether bonds in the structure, and the absorption peak at 820 cm^-1^ was attributed to the C-H out-of-plane bending vibration of the B-ring aromatic ring para substituted structure.

**Figure 10.**
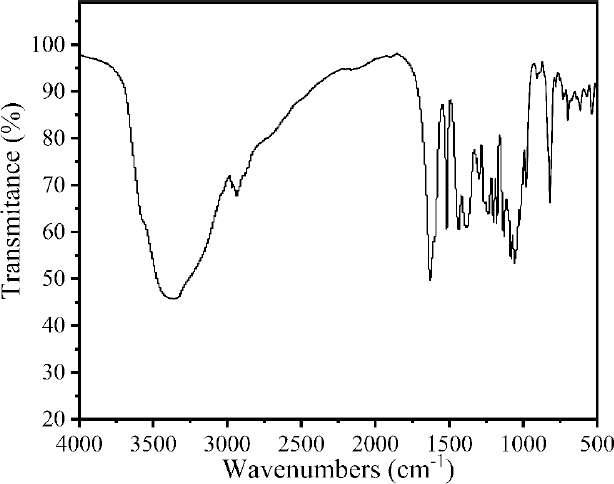
Fourier transform infrared spectrum of naringin dihydrochalcone

### 3.3.2 NMR spectrum analysis

The NMR analysis of our refined naringin dihydrochalcone products is shown in Figure 11-14. As shown in Figure 15-18, the ^1^H NMR and ^13^C NMR spectra of our refined naringin dihydrochalcone products is highly consistent with those of the standard.

Naringin dihydrochalcone sample

^1^H NMR (500 MHz, Methanol-d4) δ 7.06 (d, J = 8.2 Hz, 2H), 6.72 (d, J = 8.0 Hz, 2H), 6.07 (s, 2H), 5.29 (d, J = 1.7 Hz, 1H), 5.04 (d, J = 7.5 Hz, 1H), 4.02 – 3.87 (m, 3H), 3.73 (dd, J = 12.1, 5.3 Hz, 1H), 3.69 – 3.58 (m, 3H), 3.45 (dtd, J = 16.7, 9.6, 8.6, 5.3 Hz, 3H), 3.33 – 3.26 (m, 2H), 2.87 (t, J = 7.8 Hz, 2H), 1.34 (d, J = 6.3 Hz, 3H).

^13^C NMR (126 MHz, Methanol-d4) δ 205.68, 164.00, 163.20, 132.49, 128.97, 114.78, 101.08, 97.93, 94.90, 77.64, 77.51, 76.72, 72.61, 70.83, 70.76, 69.86, 68.56, 60.95, 46.10, 29.82, 16.89.

Naringin dihydrochalcone standard

^1^H NMR (500 MHz, Methanol-d4) δ 7.06 (d, J = 8.4 Hz, 2H), 6.72 (d, J = 8.2 Hz, 2H), 6.07 (d, J = 1.8 Hz, 2H), 5.29 (s, 1H), 5.04 (d, J = 7.4 Hz, 1H), 4.04 – 3.89 (m, 3H), 3.73 (dd, J = 12.1, 5.3 Hz, 1H), 3.71 – 3.58 (m, 3H), 3.51 – 3.38 (m, 3H), 3.30 (ddd, J = 8.8, 6.8, 2.3 Hz, 2H), 2.86 (t, J = 7.8 Hz, 2H), 1.34 (dd, J = 6.2, 1.6 Hz, 3H).

^13^C NMR (126 MHz, Methanol-d4) δ 205.70, 163.98, 163.18, 132.49, 128.98, 114.80, 101.07, 97.93, 94.91, 77.63, 77.51, 76.71, 72.62, 70.83, 70.76, 69.87, 68.56, 60.95, 46.10, 29.81, 16.91.

**Figure 11.**
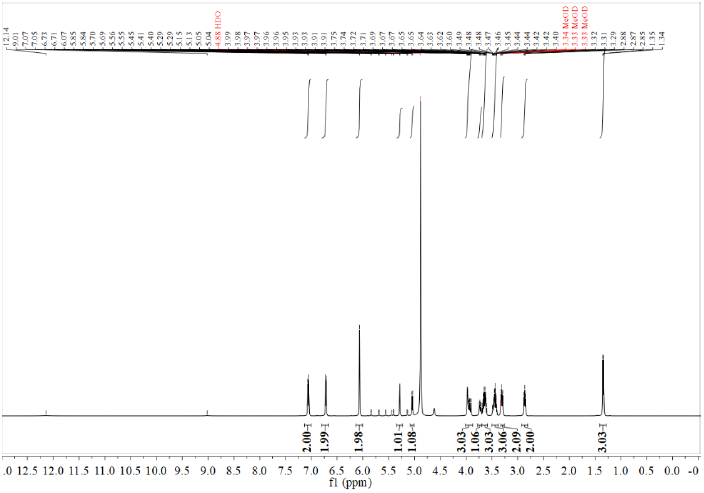
^1^HNMR spectrum of refined naringin dihydrochalcone products

**Figure 12.**
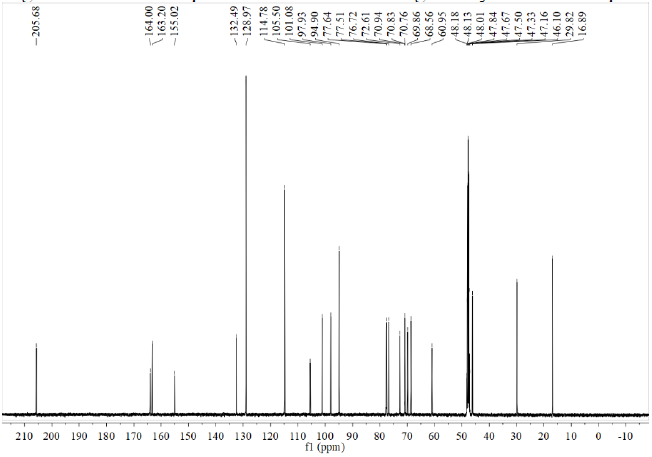
13C NMR spectrum of refined naringin dihydrochalcone products

**Figure 13.**
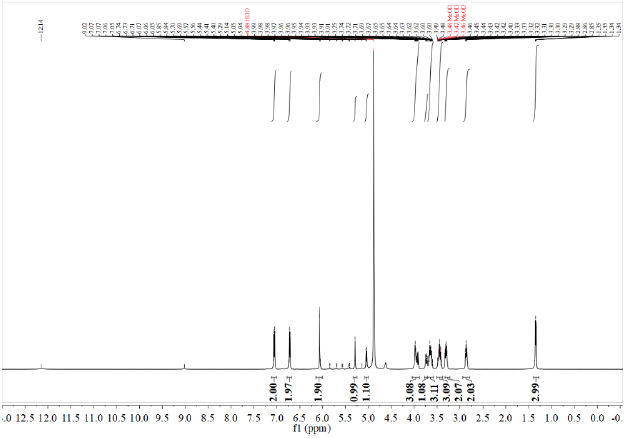
^1^H NMR spectrum of naringin dihydrochalcone standard

**Figure 14.**
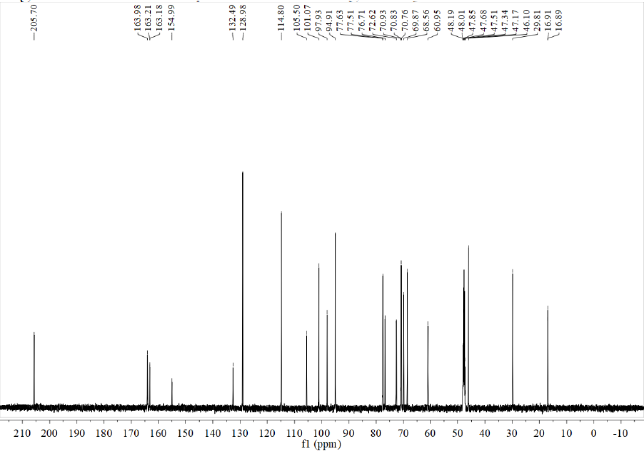
^13^C NMR spectrum of naringin dihydrochalcone standard

### 3.4 Cholate standard curve

Cholate standard curve are shown (Supplementary Materials Figure S7-8).

### 3.5 The binding ability of naringin dihydrochalcone to cholate

As seen in Figure15, with an increase in naringin dihydrochalcone concentration, the binding rate of sodium glycocholic acid and sodium taurocholate increase significantly (*p*<0.05). When the concentration of naringin was 0.5 mg/mL, the binding rate of naringin dihydrochalcone and sodium glycocholic acid was 65.29%, and that of naringin dihydrochalcone and sodium taurocholate was 37.84%, indicating that the binding capacity of naringin dihydrochalcone and sodium glycocholic acid was stronger than that of sodium taurocholate. This may be related to the spatial structure of naringin dihydrochalcone molecules, and the specific reasons need to be further confirmed.Therefore, the higher the binding rate of naringin dihydrochalcone with sodium glycocholic acid and sodium taurocholate, the higher the binding capacity of sodium cholate, the stronger the blood lipid-lowering function, and the more significant the blood lipid-lowering effect.

**Figure 15.**
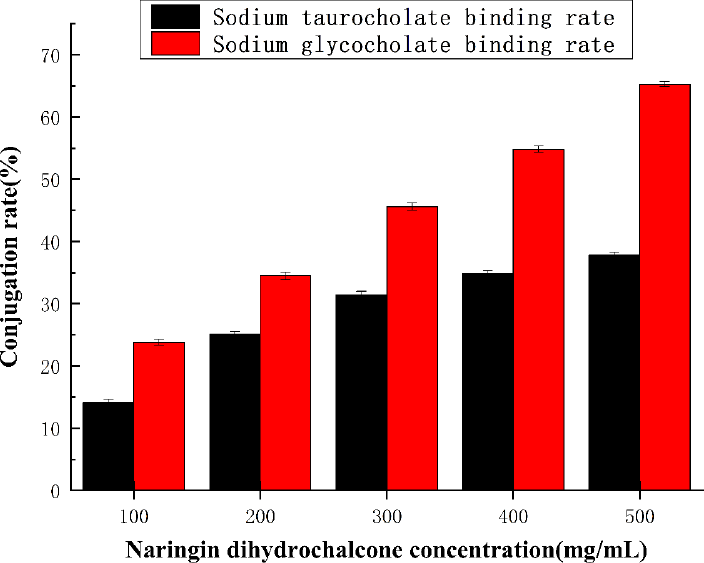
Binding capacity of naringin dihydrochalcone to cholate

### 3.6 MTT experimental results

The inhibitory effects of nine concentrations of naringin dihydrochalcone on HepG2 liver cancer cells are shown in Figure 16, which showed that at mass concentrations in the 6.25–1600μg/mL range, the survival rate of the HepG2 cells > 99.00%. This indicated that naringin dihydrochalcone within this concentration range did not exert any cytotoxic effect on Hep G2 cells, and the differences between these different concentrations were not significant (*P*>0.05). Thus, subsequent experiments could be conducted using samples with concentration in this range.

**Figure 16.**
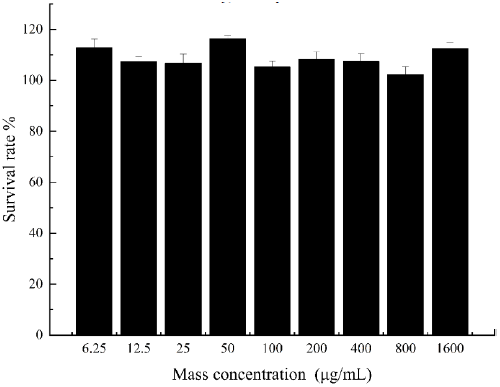
Effect of naringin dihydrochalcone on the survival of Hep G2 liver cancer cells

### 3.7 Establishment of the high-fat cell model

The construction of the high-cholesterol cell model using Hep G2 liver cancer cells is shown in Figure 17. From this figure, it is evident that when the mass concentration of cholesterol was less than 25 μg/mL, the content of cholesterol in the cells increased with increasing cholesterol concentration. However, cholesterol mass concentrations above 25 μg/mL did not result in any significant increase in cholesterol content (*P*>0.05). Further, when the mass concentration of cholesterol was 25 μg/mL, the corresponding cholesterol content was 7.41 mmol/L, higher than the normal level. This observation indicated that with a cholesterol mass concentration of 25 μg/mL, the high-cholesterol cell model was successfully constructed.

**Figure 17.**
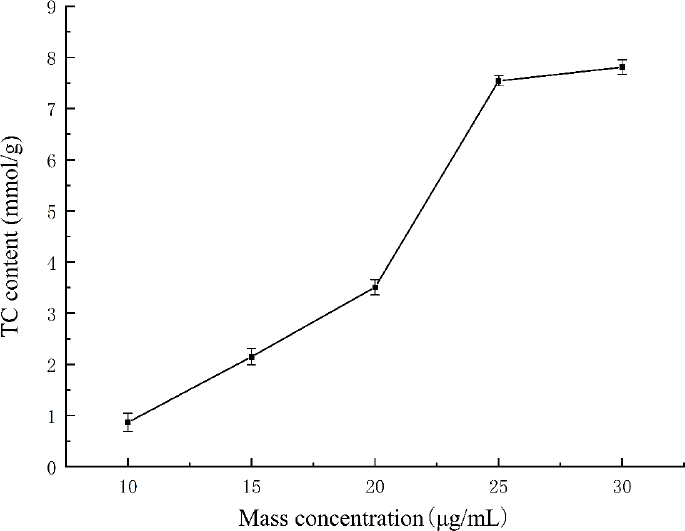
Effect of different concentrations of cholesterol on the total cholesterol content of Hep G2 cells.

### 3.8 Effects of naringin dihydrochalcone on TC, TG, LDL-C, and HDL-C levels in Hep G2 cells

Cells in the normal control and high-cholesterol model groups were treated with different concentrations (10, 20, and 40 μg/mL) of naringin dihydrochalcone to study the effects of this agent on TG and TC levels in Hep G2 cells. The changes in TG content in the cells are shown in Figure 18. Within the10–40 μg/mL concentration range, the TC and TG contents of the cells decreased with increasing naringin dihydrochalcone concentration. Further, compared with the blank group, the model group showed significantly higher TC, TG, and LDL-C levels (*P*<0.01), while the levels of HDL-C were significantly reduced (*P*<0.01). Further, compared with the model group, the low-, medium-, and high-dose naringin dihydrochalcone groups showed 20.36, 33.77, and 68.22% decreases in TC levels, 21.42, 41.78, and 55.44% decreases in TG levels, and 11.65, 20.15, and 28.71% decreases in LDL-C levels, respectively, and all these differences were statistically significant (*P*<0.01). Further, HDL-C levels increased by 33.87, 51.48, and 56.11%, respectively, and these increases were significant and dose-dependent (*P*<0.01).

**Figure 18.**
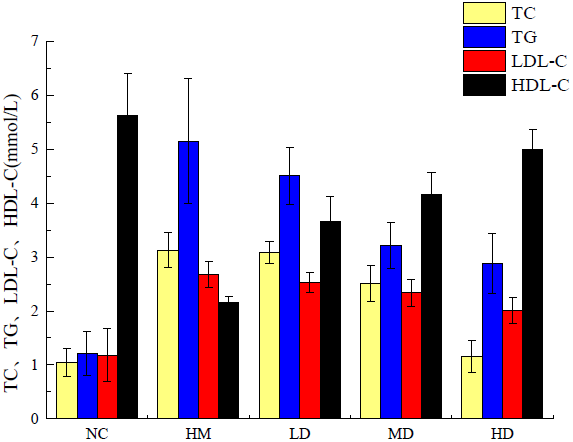
Effect of naringin dihydrochalcone on blood lipid levels in mice (Normal control, NC; Hyperlipidemia model group, HM; High dose group, HD; Media dose group, MD; Low dose group, LD)

## 4. Conclusions

In this study, we investigated optimal naringin extraction conditions. Thus, we identified extraction temperature, time, and material to liquid ratio of 67 ℃, 2.8 h, and 54:1 mL/g, respectively, as optimal for naringin extraction. Further, the yield of naringin was 32.48 mg/g, and via IR and NMR, its structure as well as that of naringin dihydrochalcone were confirmed. The determination of the lipid-lowering activity of naringin dihydrochalcone in vitro was also investigated. Specifically, we used cholesterol induction to establish a high-fat Hep G2 cell model, and compared with the model group, naringin dihydrochalcone significantly reduced intracellular TC and TG levels, indicating that it exerts a cholesterol metabolism promoting effect and can effectively prevent lipid deposition in blood vessels and the liver, while also showing a lipid-lowering effect. Cholesterol is a lipophilic substance that can bind with proteins to form lipoproteins, which dissolve in the bloodstream. Specifically, cholesterol in the LDL-C form easily adheres to blood vessel walls, resulting in an increase in blood lipid levels. However, in the HDL-C form, it can clear the bad cholesterol on the blood vessel wall and unblock blood vessels. The results of this study indicated that high-dose naringin can effectively inhibit increases and decreases in LDL-C and HDL-C levels, respectively, in a high-fat Hep G2 cell model, thus promoting cholesterol metabolism. These findings indicated that naringin dihydrochalcone has potential for application as a functional food resource for lowering blood lipid levels.

## Author Contributions

Author Contributions: L.Z. (Lei Zhang) and XL.Y. (Xiao-lei Yu) designed the study, interpreted the results, and provided solutions for experimental problems. MY.T. (Mu-yang Tian) drafted the manuscript and revised the paper. XL.Y. (Xiao-lei Yu) and SQ.L. (Si-Qi Li) performed the experiment. JW.Q. (Jing-Wen Qi) assisted with the experiment. MY.T. (Mu-yang Tian) performed data analyses and helped design the experiment. All authors discussed the results and commented on the manuscript.

## Acknowledgments

The authors thank the MOE Key Laboratory for Nonequilibrium Synthesis and Modulation of Condensed Matter, School of Physics, Xi’an Jiaotong University (Xi’an 710049, China) for their technical assistance.

